# Myosin activity drives entangled actin networks out-of-equilibrium – a quantitative approach

**DOI:** 10.64898/2026.06.08.730891

**Authors:** Nilay Çiçek, Kim V. Stöbener, Burkhard Geil, Zahra Aghaei, Andreas Janshoff

## Abstract

ATP-driven myosin II activity remodels actin networks and drives cytoskeletal matter out of thermal equilibrium, but how ATP concentration controls these dynamics remains difficult to isolate in vivo. Here, we reconstitute minimal actomyosin networks from purified components and combine passive microrheology with mean back relaxation (MBR) analysis to quantify ATP-dependent nonequilibrium fluctuations. Nonequilibrium activity is strongest at intermediate ATP concentrations (0.2-0.5 mM) and decreases at higher ATP. While single-bead van Hove distributions are approximately Gaussian, pooled distributions display apparent tails caused mainly by bead-to-bead heterogeneity rather than frequent active bursts. MBR, however, reveals clear time-irreversible dynamics by distinguishing restoring relaxation from persistent active motion. Comparing activity with network stiffness suggests a trade-off between ATP-dependent stiffening and myosin-driven remodeling. A minimal active Langevin simulation reproduces the observed MBR phenomenology, supporting a picture in which rare myosin-driven cage rearrangements generate detectable nonequilibrium signatures. These results establish MBR as a sensitive probe of active matter behavior in actomyosin networks.

**Significance Statement:** Cells operate out of equilibrium, yet the specific role of ATP concentration in driving cytoskeletal activity remains difficult to isolate in vivo. By reconstituting minimal actomyosin networks and applying passive microrheology, we directly quantify how ATP levels modulate out-of-equilibrium fluctuations. The application of mean back relaxation (MBR) analysis thereby provides a clear and broadly accessible measure of broken time-reversal symmetry that surpasses conventional analysis methods. Our results reveal an inverse relationship between ATP concentration and network dynamics, arising from different modes of myosin activity and ATP-dependent network stiffening. This work provides a quantitative framework for linking biochemical energy supply to mechanical activity in reconstituted cytoskeletal systems, offering new insights into cellular self-organization and energy-dependent regulation.

## Introduction

Living cells are inherently non-equilibrium systems that continuously consume chemical energy to carry out essential processes such as division, transport, adhesion, and migration. Many of these dynamic functions arise from the interplay between actin filaments and myosin motors (1, 2). The actin cytoskeleton therefore exemplifies this active matter paradigm in which myosin motors convert ATP into mechanical work, walking along or sliding actin filaments to generate directed motion, contractile forces, and persistent remodeling that keeps cells far from thermal equilibrium (1–9). The extent of network remodeling depends critically on network composition including crosslinker density, myosin isoform and concentration as well as ATP availability (2, 10–12). In vitro reconstituted actomyosin gels reproduce these features. Motor activity initially fluidizes networks (13) and later drives contractile cluster formation in weakly crosslinked systems (9, 14–17). Increasing myosin concentration accelerates this coarsening and produces larger clusters (14, 16). Embedded tracer beads reveal enhanced non-equilibrium fluctuations through elevated mean squared displacements and non-Gaussian statistics (15, 18). A central challenge is, however, finding means to distinguish between network heterogeneity and non-thermal out-of-equilibrium fluctuations responsible for emergent behavior of active systems. Also, a quantitative measure of out-of-equilibrium processes that can conveniently be applied to real world problems is lacking. This quantification traditionally requires invasive perturbations or combined active and passive microrheology using optical tweezers, which can be challenging in fragile biological materials and throughput is typically sparse (19). Recently, however, mean back relaxation (MBR) was introduced as a non-invasive alternative based solely on passive particle tracking (20, 21). MBR is a multi-point observable correlating particle positions at three time points, measuring the ratio of future to past displacements over defined intervals. In thermal equilibrium with detailed balance, MBR converges to exactly 0.5 and any deviation in stationary systems marks broken detailed balance and active energy injection. This makes MBR particularly powerful for detecting non-equilibrium dynamics in biological systems rendering the combination of active and passive measurements of probes obsolete. Here, we examine the impact of ATP-concentration on the out-of-equilibrium properties of the actin cytoskeleton in the presence of myosin motors. While ATP availability critically regulates actomyosin activity, in living cells ATP fuels many competing processes, obscuring its specific effect on cytoskeletal mechanics (22). To isolate this relationship, we employ a minimal reconstituted actomyosin system with precise control over ATP concentration. Combining passive microrheology with MBR analysis, we systematically investigate how biochemical fuel availability drives these networks out of equilibrium, establishing a quantitative link between ATP concentration and detailed-balance breaking in active cytoskeletal matter. We find that individual bead trajectories obey Gaussian statistics in van Hove plots, essentially masking the activity reveal by MBR analysis. Larger ATP concentration leads to slight increase in network stiffness and concomitantly to a decline in excess energy.

## Results

Internal dynamics and mechanics of the reconstituted actin and actomyosin networks were investigated by means of passive video particle tracking as illustrated in Figure 1. The representative confocal images of 24 μM muscle actin networks without and with 0.48 μM myosin II at 40 minutes of sample age shown on the upper right side of the figure display significant structural differences between actin and actomyosin networks. The entangled actin network in the absence of myosin appears homogeneous lacking large structures (left), while the actomyosin network eventually forms contractile foci with stretched filaments between them (right), demonstrating myosin-driven remodeling through crosslinking and contraction. Below the confocal images, exemplary trajectories for a bead with a diameter of 2 μm embedded in a neat, entangled actin network (blue, left) and for a bead incorporated in an active actomyosin network (red, right) are shown, respectively. While the bead in the non-contractile actin network shows spatially confined motion covering merely a radius of about 40 nm without showing any preferred directionality of the movement, the bead in the actomyosin network explores a larger space with a radius of about 1000 nm by traveling between distant positions while still performing Brownian motion. These qualitative differences in the trajectories demonstrate that the presence of myosin motor proteins has a significant impact on the displacement pattern of the probe particles.

**Figure 1.**
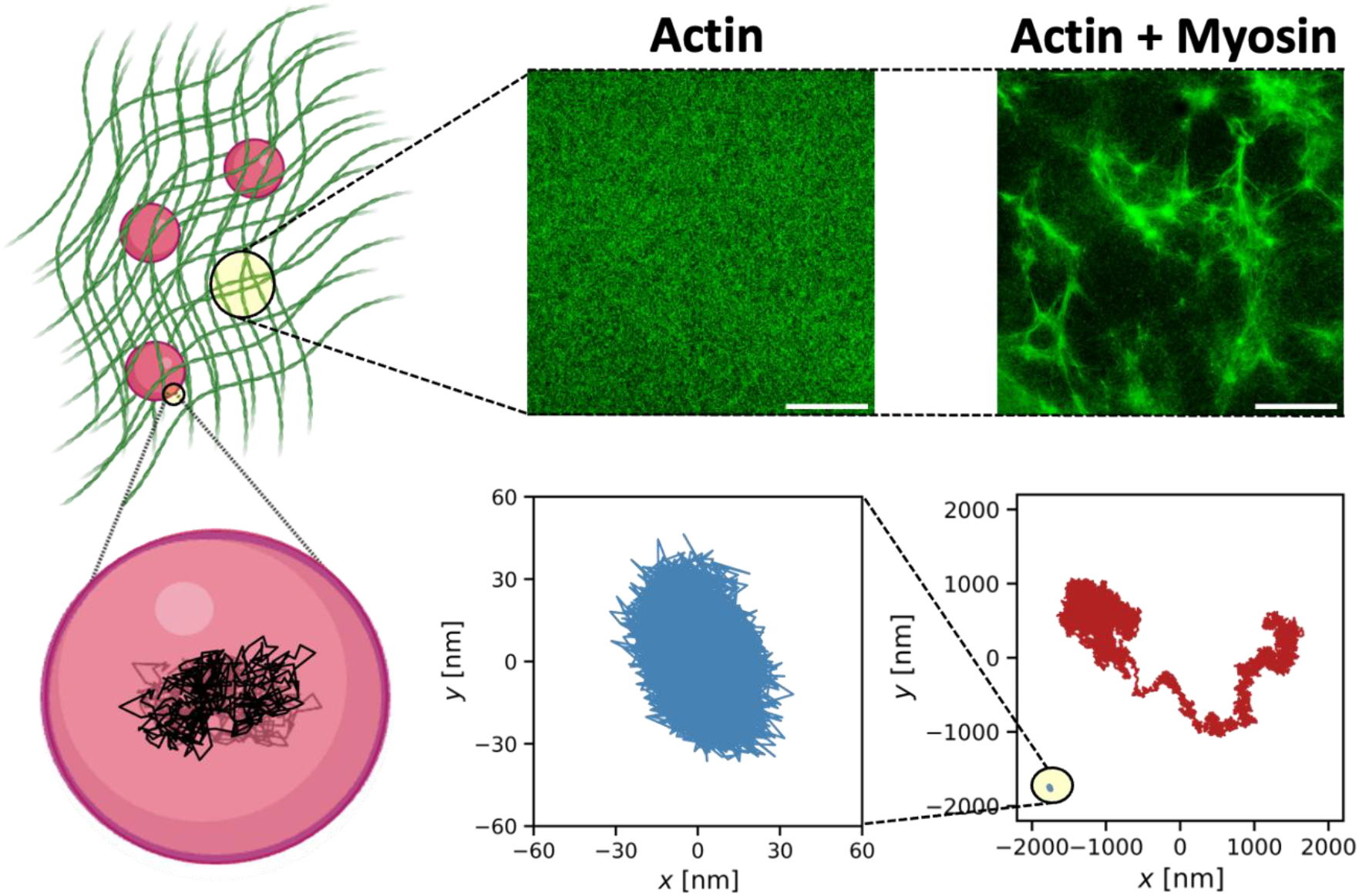
Video Particle Tracking of microparticles embedded in an actin mesh. Shown on the left is a schematic illustration of video particle tracking showing beads larger than the mesh size of the actin network and a corresponding trajectory. Exemplary confocal microscopy images of actin and actomyosin networks at a sample age of 40 minutes are shown on the upper right. The muscle actin concentration is 24 μM for both samples. 10 % of actin are ATTO-488 labelled. The myosin concentration is 0.48 μM. The scale bars represent 50 μm. The lower right panel shows characteristic trajectories of a single bead (diameter: 2 μm) embedded in an entangled actin network (blue) and in an actomyosin network (red) plotted in the same scale and a zoom on the bead trajectory in the entangled actin network on the left. Trajectories were recorded for 180 s at a rate of 138 Hz.

The question arises whether this can be explained by fluidization or active matter physics. First, we classified the data we obtained for all tested conditions. We plotted them according to their radial spread ⟨*dR*⟩/⟨*dr*⟩ and their mean step size between frames (Figure 2a). This plot permits us to identify trajectories that exhibit long distance displacements versus those which display confined motion. Each individual point in the scatter plot indicates a single trajectory of a bead either embedded in a “passive” entangled actin network (blue) or in an “active” actomyosin network (red). The plot shows a mixed population of trajectories from “active” and “passive” samples at low radial spread and a pure “active” population having higher radial spread. Among our data, we observed different types of trajectories as illustrated by the characteristic trajectories I, II, III and IV below the scatter plots (Figure 2b). Type I and II trajectories are typically found in the mixed population of both scatter plots and show a rather symmetric spread around their center of mass (Figure 2b). Type III trajectories are typical for the “active” population and display more directionality and higher spread, while type IV trajectories illustrate an unusual case, where persistent trends dominate (Figure 2b). Below the characteristic trajectory plots, their respective MSD graphs are shown (Figure 2b). While the MSDs of trajectory type I and II have a slope far below 1 and approach a plateau indicating confinement (23), the MSDs of trajectory type III and IV increase only slightly at first with increasing lag time followed by a steep increase with a slope of ∼ 1 without reaching a plateau (Figure 2b). Thereby, the first regime accounts for the fluctuations at early lag times, and the second regime accounts for the flow of the entire network. While MSDs give information on the type of bead motion and physical constraints, it provides no information on whether the system is out of equilibrium. A common analysis to demonstrate activity in terms of non-Gaussian behavior is the Van Hove distribution (15, 18). However, non-Gaussian tails in the ensemble Van Hove distribution of our actomyosin networks arise solely from network heterogeneity as Van Hove distributions of individual beads are pure Gaussians with different widths (Figure S1). Furthermore, Van Hove distributions do not directly quantify the degree of detailed-balance breaking or provide a direct measure of how far the system operates from thermal equilibrium. Hence, to quantitatively assess network dynamics, we employed mean back relaxation (MBR) analysis, a recently developed method that quantifies deviations from thermal equilibrium through three-point correlation statistics (20, 21) thus it is sensitive to non-Markovian or hidden-Markovian effects. Unlike traditional two-point observables such as MSD, MBR correlates particle positions at three sequential time points and measures how a particle’s future displacement relates to its past trajectory (20, 21). The mean back relaxation (MBR) quantifies how a particle’s future displacement is related to its preceding displacement. Consider three positions along a trajectory: the particle position *x*(−*τ*) at a time −*τ* before a reference time, the reference position *x*(0) = *x*_0_, and the future position *x*(*t*) = *x*. We define the preceding displacement *d* = *x*_0_ − *x*(−*τ*) and the subsequent displacement *b* = *x*(*t*) − *x*_0_. The MBR measures the expected normalized relaxation displacement,

**Figure 2.**
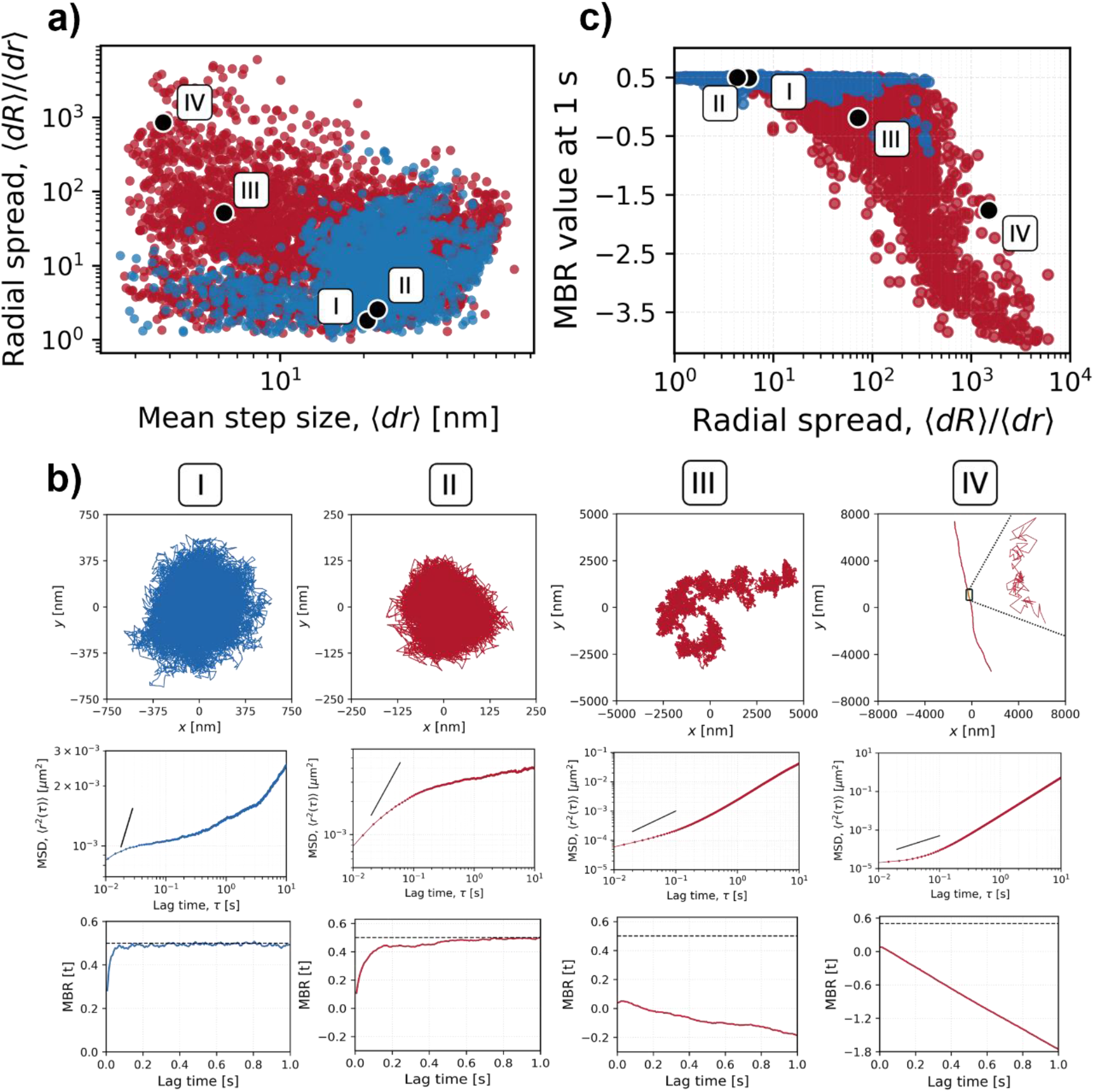
Classification of bead trajectories. a) Feature-space mapping of trajectories based on their mean step size ⟨*dr*⟩ between consecutive frames and radial spread to step size ratio ⟨*dR*⟩/⟨*dr*⟩ with characteristic trajectories marked. Each datapoint in the scatter plot represents a single trajectory of a bead embedded in actin (blue) or actomyosin networks (red) of all tested conditions. b) The most commonly observed characteristic trajectories were classified into four categories based on their defining features. According to this classification, two-dimensional trajectories, their corresponding MSD and MBR curves are shown. The black line in MSD curves represents a slope of 1. c) Feature-space mapping of trajectories based on their radial spread ⟨*dR*⟩/⟨*dr*⟩ and their MBR value at a lag time of 1 s with the same characteristic trajectories marked. Each datapoint in the scatter plot represents a single trajectory of a bead embedded in actin (blue) or actomyosin networks (red) of all tested conditions. Trajectories were measured at a sample age of 20 minutes. Actin concentration: 24 μM, myosin concentration: 0.48 μM. The analysis was conducted with a total of 6179 trajectories.

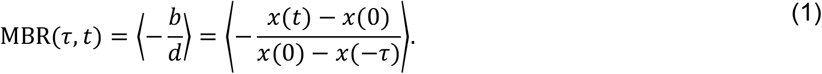

With this sign convention, positive MBR values indicate back-relaxation: after a preceding displacement *d*, the particle tends to move in the opposite direction. Negative MBR values indicate persistence: the subsequent displacement *b* tends to have the same sign as the preceding displacement *d*. Equivalently, the MBR can be written in terms of the three-point probability density *W*_3_(*x, t*; *x*_0_, 0; *x*_0_ − *d*, −*τ*) as

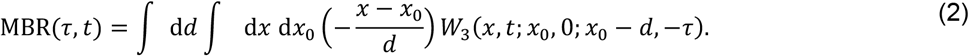

Here, the three-point density weights all triplets of trajectory points (*x*(−*τ*), *x*(0), *x*(*t*)) according to their probability of occurrence (20). Over time the MBR generally saturates and approaches a plateau. In systems exhibiting detailed balance and thermal equilibrium, MBR converges to a value of 0.5 for long observation times, while deviations from this value indicate broken detailed balance and non-equilibrium activity (20). Furthermore, MBR has been shown to correlate linearly with the effective energy amplitude *E*_0_, providing a direct link between passive trajectory measurements and the energy scale of active fluctuations in the system (Equation 3).

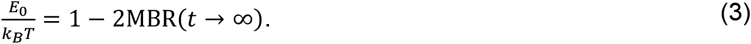

Figure 2c shows the measured trajectories plotted according to their MBR plateau heights at an arbitrary chosen lag time of 1 s and their radial spread ⟨*dR*⟩/⟨*dr*⟩. Again, a mixed population and a purely active population occur. The mixed population of trajectories from “active” and “passive” samples shows a rather low radial spread and MBR values around 0.5 indicating equilibrium. This suggests that actomyosin samples do not always display activity as out-of-equilibrium properties (Figure 2c). The pure “active” population consisting only of trajectories of beads embedded in actomyosin networks shows strong deviations from an MBR value of 0.5 and trajectories often display higher radial spread. The MBR graphs of the characteristic trajectories of type I, II, III and IV are shown in the last row of Figure 2b. For type I and II trajectories, MBR approaches a value of 0.5, indicating detailed balance. For type III and IV trajectories, MBR lies well below 0.5. Especially for type IV trajectories exhibiting strong persistent motion, MBR decreases linearly with increasing lag time indicating continuous memory loss caused by the network flow (Figure 2b). Since MBR theory is restricted to confined systems (20) and network flow diminishes bead confinement, we decided to detrend our trajectories as illustrated in Figure S2 and thereby focus on short timescale fluctuations rather than flow events for the proceeding comparison of networks with different ATP concentrations.

Figure 3 displays fluorescence microscopy images of actin and actomyosin networks at increasing ATP concentrations (0.5 mM, 1.0 mM, 1.2 mM, 2.0 mM). Actin networks remain homogeneous up to 1.2 mM ATP but cluster at higher concentrations probably due to electrostatic interactions between negatively charged ATP and actin filaments, as well as enhanced polymerization (Figure 3, Figure S3). With myosin present, networks form contractile foci at all ATP concentrations already at 20 minutes of sample age with diameters ranging from a few microns at 0.5 mM ATP up to around 30 μm at 2 mM ATP. Above 1 mM ATP, clusters also show higher fluorescence intensity and fewer connecting filaments, indicating stronger coarsening (Figure 3, Figure S3).

**Figure 3.**
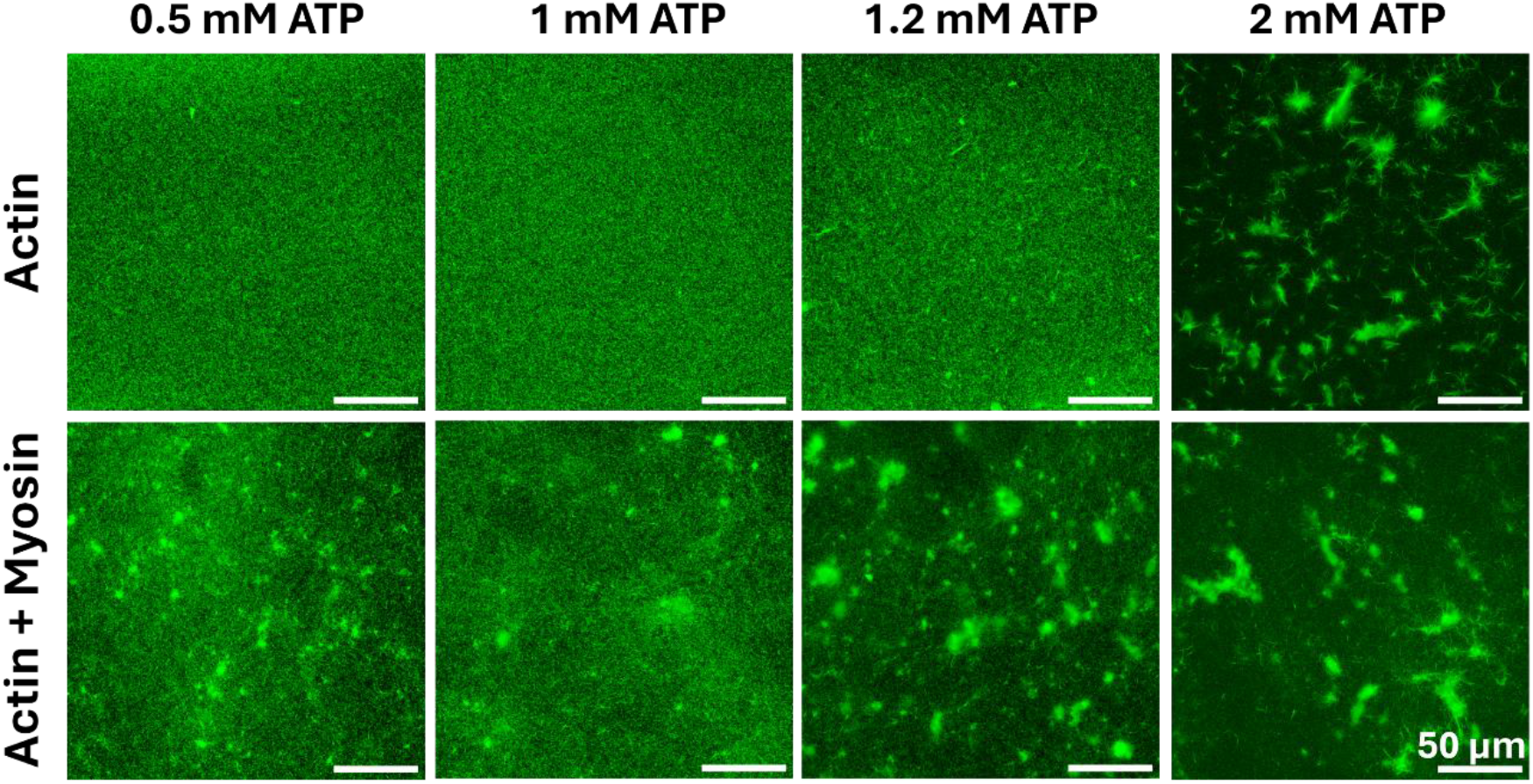
Confocal fluorescence images of actin and actomyosin networks polymerized at increasing ATP concentrations (from left to right). The images were taken 20 minutes after sample preparation. Histogram equalization was applied in Fiji to enhance contrast (54). Actin concentration: 24 μM with 10% being ATTO-488 labelled actin; myosin concentration: 0.48 μM. The scale bars represent 50 μm.

Figure 4a shows the MSD of beads embedded in entangled actin networks with increasing ATP concentrations at 20 minutes of sample age. For all ATP concentrations, the MSD curves display a power law exponent α lower than 1 (subdiffusive) and eventually reach a mean plateau corresponding to a cage size of 100 nm for 2 mM ATP and 220 nm for 1 mM ATP at a lag time of about 10 seconds (Figure 4a), which indicates confinement (23). The power law exponent α obtained as the slope of the MSD in logarithmic scale measured between lag times *τ* = 0.1 s and *τ* = 1 s is low for the highest ATP concentration (2 mM) (Figure 4c). This indicates a higher network stiffness, which we confirmed by converting the MSDs of those passive samples into their corresponding rheological spectra using a standard procedure based on the generalized Stokes– Einstein equation (GSE), which links microscopic bead fluctuations to the material’s viscoelastic response (24, 25). From the resulting frequency dependent shear modulus, we then identified the plateau modulus *G*0 by examining the region around 0.1 Hz, where the modulus becomes relatively frequency independent and reflects the network’s elastic stiffness (26, 27). The plateau modulus *G*0 ranges from 0.1 to 1 Pa and slightly increases with increasing ATP concentration at all sample ages (Figure S4). For active samples, the GSE is not fulfilled, and G moduli should not be calculated. In the presence of myosin, the networks become more fluid as indicated by the increased α values (less subdiffusive on shorter time scales) (Figure 4b/c). The highest power law exponents α are observed for actomyosin networks with 0.2 and 0.5 mM ATP (Figure 4c). However, plotting the ratio of the MSD scaling exponent α of the particles in the actomyosin network and in the corresponding neat actin network without myosin shows almost equally strong fluidization effects by myosin on networks with 1.2 mM and 2 mM ATP (Figure 4c). Taken together, the embedded beads experience larger displacements in actomyosin gels, are faster, and their movement is less restricted.

**Figure 4.**
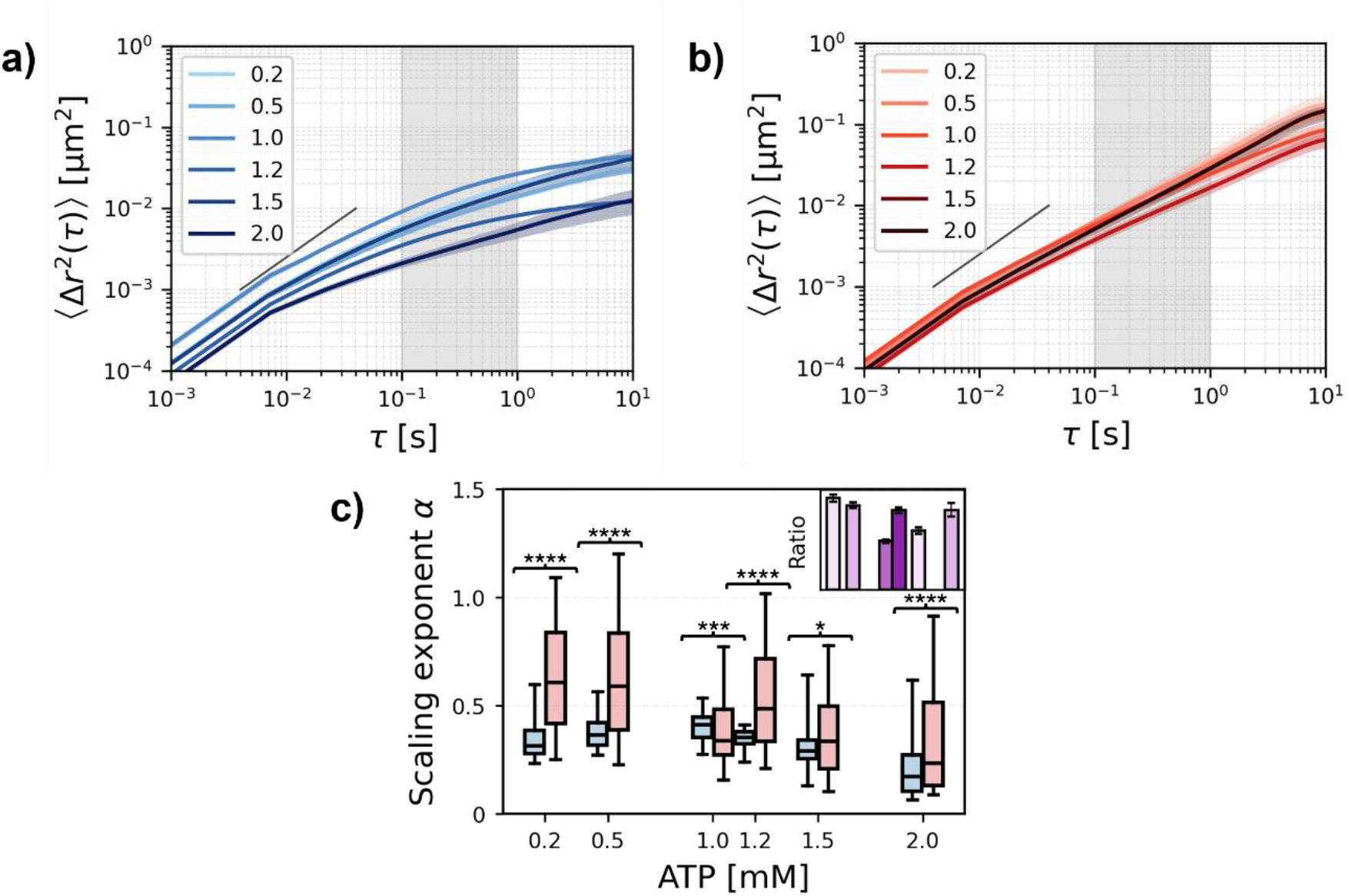
Mean squared displacements (MSDs) computed from detrended two-dimensional trajectories of probe particles embedded in actin and actomyosin networks at all tested ATP concentrations. a) Ensemble-averaged MSD curves of beads embedded in actin networks with different ATP concentrations (0.2 mM, 0.5 mM, 1 mM, 1.2 mM, 1.5 mM, 2 mM) at a sample age of 20 minutes. Underlying trajectories were detrended to remove drift and network flow. MSDs are plotted only up to *τ* = 10 s to focus on small time scales. b) Similar to a, but for actomyosin networks. Panels a and b show the mean curves, computed by averaging over all MSDs of all trials. The shaded region around each curve corresponds to the mean ± SEM. The black line shows a slope of 1. c) Mean slope α between *τ* = 0.1 s and *τ* = 1 s. Boxplots show median, interquartile range (IQR), and 1.5×IQR whiskers. To highlight the change in probe-particle diffusion, ratios between the values for actomyosin and respective actin networks are shown in the upper right corner with their propagated errors. Significance was assessed using a two-sided Mann–Whitney U test comparing MSD slopes and plateau heights for actomyosin networks and the respective networks without myosin. *, **, ***, and **** indicate p < 0.05, 0.01, 0.001, and 0.0001, respectively. The respective standard errors and p-values are listed in Table S2. The number of trials and individual probes for each sample condition are listed in Table S1. Actin concentration: 24 μM, myosin concentration: 0.48 μM.

Having established that ATP concentration profoundly affects network structure and bead dynamics, a fundamental question emerges: to what extent are these networks driven out of equilibrium, and how does this non-equilibrium character depend on ATP availability? Therefore, we carried out MBR analysis of the bead trajectories as a function of ATP concentration. The ensemble averaged results of MBR analysis are shown in Figure 5. Entangled actin networks without myosin exhibit MBR plateaus at approximately 0.5, confirming detailed balance dominated by thermal fluctuations, with corresponding effective energies near zero (Figure 5a/c). In contrast, actomyosin networks exhibit MBR values below 0.5, indicating non-equilibrium activity driven by motor-generated forces (Figure 5b). This deviation from 0.5 quantifies the extent to which myosin activity dominates over thermal fluctuations and reflects the temporal asymmetry in bead trajectories: past displacements actively influence future motion due to myosin-driven network reorganization, violating the time-reversibility expected in thermal equilibrium. The corresponding effective energies are well above zero, confirming active network behavior (Figure 5c). The low ATP concentrations of 0.2 mM and 0.5 mM produce exceptionally high activity that surpasses that of all other conditions (Figure 5c).

**Figure 5.**
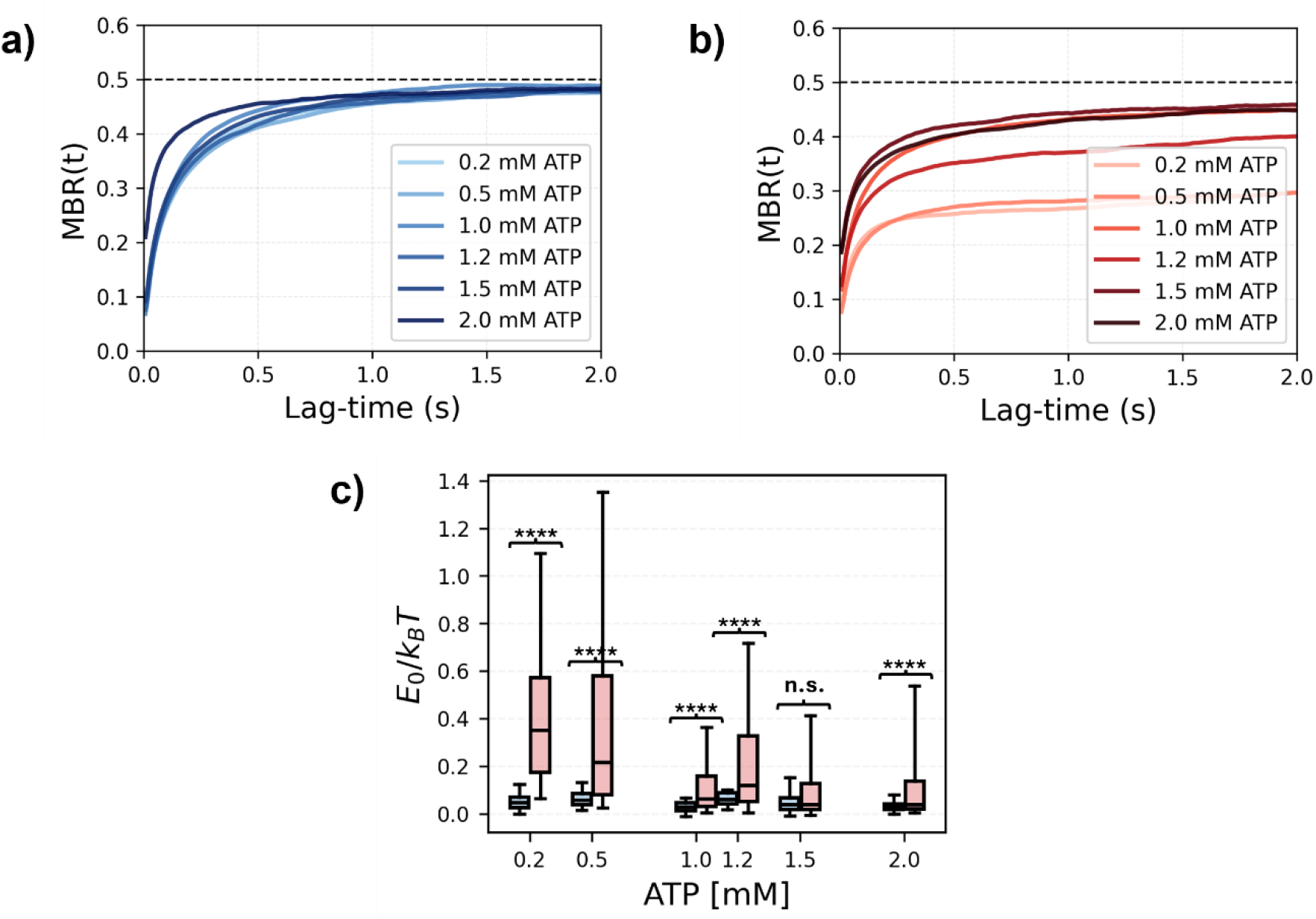
Mean back relaxation (MBR) and corresponding effective energies *E*_0_/*k*_*B*_*T* of actin and actomyosin networks at a sample age of 20 minutes. a) Ensemble averaged MBR curves of detrended trajectories of beads embedded in actin networks; each curve corresponds to one ATP concentration. The underlying trajectories were detrended using a moving average filter to remove ballistic flow and focus on small scale fluctuations. The color gradients from dark to light represent a progressive decrease in the ATP concentration (2 mM, 1.5 mM, 1.2 mM, 1 mM, 0.5 mM, 0.2 mM) b) MBR curves of actomyosin networks; each curve corresponds to one ATP concentration. The color gradients from dark to light represent a progressive decrease in the ATP concentration (2 mM, 1.5 mM, 1.2 mM, 1 mM, 0.5 mM, 0.2 mM) c) Effective energies calculated from long-term MBR values (*τ* > 1 s) for the quantification of network activity. All pairwise comparisons between conditions were performed using two–sided Mann–Whitney U test. Significance levels were indicated as p<0.05 (*), p<0.01 (**), p<0.001 (***) and p<0.0001 (****). All p-values and sample statistics can be found in Table S1 and Table S3. Actin concentration: 24 μM, myosin concentration: 0.48 μM, trajectories recorded 20 minutes after the polymerization has started.

The picture emerging from the experimental data is that of a thermally fluctuating bead embedded in a complex viscoelastic environment, whose passive dynamics are reminiscent of a generalized Langevin equation, but which is additionally subject to nonequilibrium random kicks generated by myosin motor activity. These bursts give rise to observable activity, enhanced fluidity, and deviations from purely passive viscoelastic relaxation. We therefore sought to translate these physical insights into a minimal simulation framework to determine how model parameters affect observables such as the MSD, van Hove displacement distributions, and, most importantly, MBR. In this simplified active hidden-cage model, the bead is treated as an overdamped particle with position *x*(*t*), elastically coupled to a hidden moving cage center *q*(*t*) and to a viscoelastic memory coordinate y(*t*). Active bursts do not displace the bead directly; instead, they generate random jumps in the hidden cage velocity, causing the cage center *q*(*t*) to move persistently over a finite time. The bead subsequently follows this active cage motion through the elastic coupling *k*xq, while the viscoelastic coordinate *y*(*t*) provides delayed restoring memory. Figure 6 summarizes the dynamics of beads embedded in passive and active hidden-state networks. The passive trajectory (Figure 6a) is compact and remains confined close to the local cage, whereas the active trajectory (Figure 6b) explores a much larger region of space. This reflects the central physical mechanism of the model: in the active case, intermittent motor-like bursts move the hidden cage, and the bead is transported by its elastic coupling to this moving network environment. Thus, the active trajectory represents a bead embedded in a contractile actomyosin-like network whose local mesh is intermittently displaced by motor-generated stresses. The van Hove distributions (Figure 6c/d) provide a more detailed view of the displacement statistics. At the shorter lag time, *τ* = 1 (dimensionless, see methods), the passive and active distributions are both close to Gaussian, with only very weak deviations in the tails. This is consistent with the fact that, over short times, displacements are still dominated by local thermal fluctuations and linear cage response. At the longer lag time, *τ* = 5, the active distribution broadens substantially relative to the passive distribution. The active data remains approximately Gaussian despite the burst process, indicating that many independent cage and bead fluctuations contribute to the displacement statistics. Thus, activity does not produce purely jump-dominated displacement statistics; rather, it broadens an otherwise near-Gaussian displacement distribution and adds moderate tailing. This is exactly what we observe in our experiments where the tailing of pooled distributions arises from heterogeneity rather than large jumps (Figure S1).

**Figure 6.**
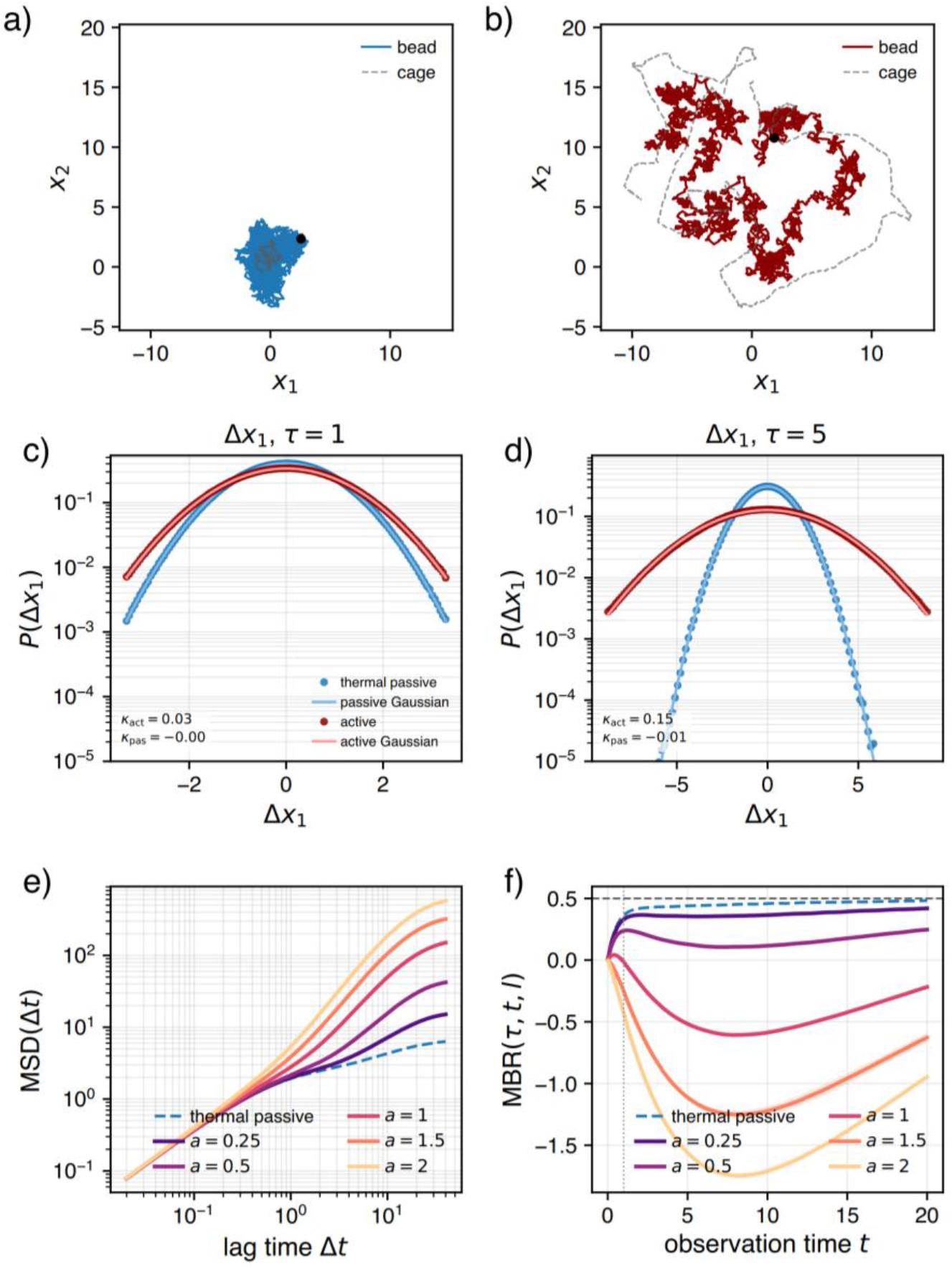
Brownian simulations of particles in an active, viscoelastic environment. (a,b) Representative trajectories in the thermal passive reference and active network. Solid curves show the observed bead position and dashed, gray curves show the hidden cage trajectory. Passive trajectories remain more confined, whereas active trajectories explore a larger region due to burst-driven cage motion. (c,d) Seed-pooled van Hove distributions of *Δx*_1_ at lag times *τ* = 1 and *τ* = 5, computed from the same passive and active baseline simulations. Gaussian fits with matched mean and variance are shown as solid curves. Despite intermittent bursts, the distributions remain close to Gaussian in the central region (see kurtosis values as inserts), while activity broadens the distribution and enhances the tails, especially at longer lag time. (e) MSD activity sweep. Increasing the activity multiplier *a*, which rescales burst amplitudes, systematically enhances long-time bead spreading. (f) MBR activity sweep. Passive trajectories show positive back relaxation, whereas increasing activity suppresses MBR and can make it negative, indicating persistent cage-driven motion rather than passive recoil. Shading denotes ±1 standard deviation across five independent random seeds. Van Hove distributions are pooled across seeds. MBR parameters are *τ* = 1.0 and 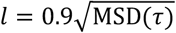. The chosen MBR conditioning time is comparable to the burst waiting time and shorter than the cage persistence time, making it sensitive to active cage rearrangements.

Similarly to the experiments shown in Figure 4, the MSD activity sweep shows that increasing burst amplitude strongly enhances longtime bead transport (Figure 6e). At short lag times, the curves remain relatively close because the bead initially responds to local thermal fluctuations and elastic confinement. At longer lag times, the curves separate systematically with activity (burst amplitude or frequency). This separation occurs once the cage has had time to move persistently after burst events and transmit this motion to the bead. The relevant time scales are therefore the burst waiting time 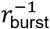, the cage persistence time *τ*_*ν*_, and the observation lag. For lags shorter than these scales, the dynamics appear more passive; for lags comparable to or longer than them, active cage transport dominates. The MBR reveals a complementary aspect of dynamics. In the passive reference, large displacements are followed by positive back relaxation, consistent with elastic recoil inside a confining cage. In contrast, increasing activity strongly suppresses the MBR and can drive it below zero over intermediate observation times. A negative MBR means that, after a selected displacement, the bead tends to continue in the same direction rather than relax back. This behavior is expected when the displacement is caused not by a passive fluctuation away from a fixed cage, but by active motion of the cage itself. The MBR is therefore a sensitive diagnostic for deviations from passive equilibrium-like behavior. While the MSD detects enhanced spreading, it cannot by itself distinguish between increased effective temperature, persistent transport, or intermittent cage rearrangements. The MBR directly probes whether a displacement is followed by recoil, as expected for a passive elastic cage, or by continued motion, as expected for active non-equilibrium driving. In the active simulations, the MBR can first decrease and later rise back toward approximately 0.5. This behavior is not contradictory and does not necessarily indicate equilibrium. The initial decrease reflects the persistence of active motion: a large displacement *d* is often caused by a recent active burst in the hidden cage velocity *ν*. Since *ν* relaxes only on the timescale *τ*_*ν*_, the cage and bead may continue to move in the same direction, so *b*/*d* > 0 and therefore −*b*/*d* < 0. A decrease of the MBR thus signals forward persistence. At longer times the active velocity memory decays. Once the system has lost memory of the selected displacement, the future position *x*(*s* + t) becomes approximately independent of *d* (*s* is an arbitrary reference time along the trajectory). For a stationary centered process (Equation 4), the conditional average of the future position, given that the previous displacement had value of *d* converges

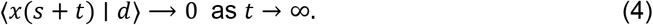

For a stationary Gaussian-like process, conditioning on *d* = *x*(*s*) − *x*(*s* − *τ*) implies approximately

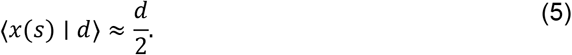

Therefore,

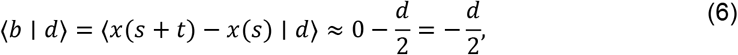

and substituting into (2) gives

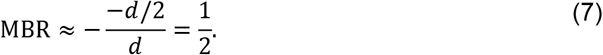

Hence, the long-time value 0.5 arises naturally for any stationary confined process once the memory of the selected displacement has decayed. This is why the MBR of the active simulation first goes down and up again with time (Figure 6f).

In the present model, the simultaneous observation of enhanced MSD, broadened van Hove distributions but with Gaussian density, and reduced or negative MBR supports the interpretation that active actomyosin-like cage rearrangements drive bead motion away from passive confined dynamics. These findings go hand in hand with our experimental observations and analysis hence the model is suitable in a sense to capture some of the essential physics of our system in which myosin motors generate bursts driving the system out of equilibrium. This is barely visible in van Hove plots showing mainly Gaussian statistics albeit broadened, while being most prominent in MBR traces. Importantly, when enhancing the burst amplitude (*a*) the MBR clearly deviates more strongly from the expected value of 0.5 indicative of being farther from equilibrium.

## Discussion

In this work, we employed passive microrheology and stochastic simulations to quantify both the mechanical properties and active dynamics of reconstituted actin and actomyosin networks as a function of ATP concentration. The goal was to assess how far from equilibrium these networks can be driven depending on composition and ATP availability. By combining multiple analytical approaches, such as mean squared displacement analysis, Van Hove distributions and mean back relaxation, we systematically characterized how myosin and ATP modulate the interplay between network stiffness and myosin-driven activity.

Mean squared displacement analysis provides a first, accessible venue into how myosin activity alters the microscopic dynamics of the actin network as a function of added ATP. In passive actin networks, MSD curves are subdiffusive across all ATP concentrations and approach a well-defined plateau at lag times of approximately 10 seconds, consistent with beads confined within a viscoelastic mesh with a mesh size between 100 and 220 nm for the tested conditions (Figure 4a). This is in accordance with previous findings from others (23). The maximal MSD of neat actin samples decreases slightly with increasing ATP concentration, reflecting enhanced network stiffening at higher ATP levels as also confirmed by the slightly increasing plateau moduli G_0_ (Figure 4a, Figure). ATP and its associated Mg^2+^ ions can mediate electrostatic interactions with the negatively charged actin filaments (28, 29). Divalent cations like Mg^2+^ are known to induce actin bundling through counterion condensation that screens electrostatic repulsion between filaments (30). Additionally, the negative charges of ATP molecules themselves may contribute to inter-filament bridging in the presence of Mg^2+^. These factors might lead to the increased network stiffness at higher ATP concentrations.as well as to the myosin-independent clustering observed at high concentrations (Figure 3; Figure S3) (31). The addition of myosin II fundamentally changes this picture: the MSD slope increases substantially, indicating that myosin fluidizes the network and effectively enlarges or abolishes the confining cage experienced by the probe particles (Figure 4b). This motor-driven fluidization has already been shown in previous studies (18, 32, 33). However, while MSD analysis sensitively captures changes in bead mobility and network mechanics, it cannot distinguish whether enhanced fluctuations originate from genuine non-equilibrium, energy-consuming processes or from passive structural heterogeneity of the network (20, 21). An elevated MSD slope in an actomyosin network relative to its passive counterpart is suggestive of active remodeling, but this difference could in principle also arise from myosin-induced changes in mesh architecture i. e. fluidization or bead–filament coupling rather than from broken detailed balance. MSD analysis thus provides valuable information on network mechanics and the extent of motor-driven fluidization but does not by itself constitute a rigorous measure of how far the system operates from thermal equilibrium. While Mizuno, Koenderink, and their respective coworkers have linked non-Gaussian tails in van Hove plots of bead trajectories to active dynamics, we demonstrate that these deviations more likely stem from sample heterogeneity, as initially proposed by Weitz and coworkers (15, 34, 35). Although rare, large-amplitude jumps driven by myosin action do occasionally give rise to these tails; our MBR analysis reveals that the system is indeed active despite adhering predominantly to Gaussian statistics. Thus, the vast majority of the active fluctuations remain hidden within the Gaussian core of the distribution (Figure S1).

A key methodological advance of our study is therefore the application of MBR analysis to reconstituted cytoskeletal networks (20, 21). By correlating three consecutive particle positions, MBR measures the degree to which a particle’s future displacement is conditioned on its immediate past, a causal asymmetry that is strictly forbidden in systems obeying detailed balance and that can only arise from active, energy-consuming processes (20, 21). The convergence of MBR to exactly 0.5 in all passive actin networks, irrespective of ATP concentration, confirms thermal equilibrium and simultaneously validates that network stiffening at elevated ATP does not by itself introduce any apparent non-equilibrium signal (Figure 5). In actomyosin networks, the pronounced depression of MBR below 0.5, and the correspondingly elevated effective energies *E*_0_/*k*_*B*_*T* provide a model-independent, quantitative measure of broken detailed balance that directly reflects the energy dissipated by myosin motors into the network. The low ATP concentrations of 0.2 mM and 0.5 mM ATP yield the highest effective energy, surpassing all other tested conditions (Figure 5). This may be surprising at first as one would expect that higher levels of ‘chemical fuel’ trigger elevated activity. However, the mechanical state of an actomyosin network is governed by the interplay between motor force generation, network connectivity, and the duty ratio of myosin II filaments (36, 37). At low ATP concentrations (0.2 mM, 0.5 mM), myosin motors exhibit slower ATPase cycling rates, leading to prolonged attachment times on actin filaments (38, 39). This kinetic regime allows motors to act as persistent crosslinkers while simultaneously generating sustained contractile forces (40). The extended duty ratio also leads to a higher number of simultaneously attached motor heads, that collectively enable more efficient force transmission and coordination across the network, driving collective network rearrangements (32, 38, 39). This goes hand in hand with our simulations showing an increased departure from MBR = 0.5 with elevated burst amplitude (Figure 6f). At higher ATP concentrations, motors cycle rapidly through their mechanochemical cycle, transitioning quickly between bound and unbound states (22, 40). This rapid turnover compromises the spatial and temporal coherence of force generation, especially at high load (39, 40) and reduces the duration of individual force-transmission events, leading to a smaller net departure from 0.5 in the MBR statistic despite the higher ATP turnover rate, consistent with the reduced effective energies and with the notion that efficient force transmission requires a minimum motor attachment time (38, 39). Additionally, myosin motors become locally depleted due to network contraction and coarsening (41). Furthermore, elevated ATP concentrations promote enhanced actin polymerization and electrostatic-mediated filament clustering potentially through Mg2+-ions, progressively stiffening the network and constraining motor-driven remodeling (Figure 2, Figure S5) (29, 42). The observed behavior is consistent with theoretical predictions and experimental observations in reconstituted actomyosin gels, where mechanical activity is maximized not at saturating ATP but at intermediate motor occupancies that optimize the coupling between force generation and network deformation (32, 39, 40).

Together, these findings establish MBR as a uniquely powerful complement to distribution-based analyses: where van Hove statistics reveal *what* kind of fluctuations are present, MBR reveals *whether* those fluctuations are thermodynamically irreversible, enabling a complete and unambiguous characterization of active matter behavior from purely passive measurements. Here, we are dealing with the special case of a subdiffusive system that exhibits predominately Gaussian van Hove statistics but is still far from equilibrium as reported by the MBR. To further rationalize this, we did computer simulations like those recently reported by Knotz et al. (43). We use a hidden Markov model to mirror the viscoelastic properties of the network by a single mode (relaxation time) to keep things simple (44). This hidden-state active cage model provides a minimal phenomenological representation of bead motion in contractile actomyosin-like networks. Note that the bead is not modeled as an active particle. Instead, activity enters through the surrounding medium in which myosin-like events only perturb the local cage velocity. Consequently, the cage moves persistently, and the bead follows through elastic coupling. This mechanism is consistent with the physical picture of embedded beads reporting local network deformation and motor-driven rearrangement. The model also clarifies the relationship between different observables (Fig. S5-S7). Back relaxation is favored in stiffer and more viscous networks, while persistence of movement is mitigated. The MSD measures the magnitude of bead spreading, the van Hove distribution resolves the full displacement statistics, and the MBR tests whether selected displacements are followed by recoil. In a passive cage, displacement and recoil are strongly linked. In an active cage, the local reference position can itself move, so recoil is reduced or reversed. This makes MBR particularly useful for identifying non-equilibrium cage dynamics that may not be obvious from the MSD alone.

The physical picture that emerges is summarized in Figure 7. Sporadic myosin-driven contractile bursts inject impulses into the hidden cage velocity. Higher burst amplitude or a high burst rate mirrors an increased ATP concentration in the experiment. Because this velocity relaxes only over the finite persistence time, these impulses are converted into transient persistent motion of the local actomyosin cage. Through viscoelastic coupling to the surrounding network, this cage motion displaces the embedded bead. Although the resulting displacement statistics remain close to Gaussian, with only weak tailing in the van Hove distributions, the non-equilibrium character of the dynamics is clearly exposed by long-time deviations of the mean back relaxation, reflecting the suppression of passive recoil after large displacements. This holds for the experiment as well as the simulations that take these assumptions (Figure 7) as ingredients. Taking together the results from all applied analysis, our systematic variation of ATP concentration reveals the following picture: the relationship between ATP supply and network activity is highly inverse, arising from effects of ATP on both motor kinetics and network mechanics. At low ATP concentration the network is softer, and motors action is more concerted both giving rise to deviations of MBR from 0.5. Our comparative results are based on relative effects associated with increasing initial ATP concentrations. The ATP-dependent behavior observed in the experimental system is dependent on externally added and increasing ATP concentrations. ATP originating from the G-buffer is negligible within the final network structure and is insufficient to explain the observed increasing ATP effect. Furthermore, rather than generalizing our claims for the entire measurement period, we limit them to the early period when ATP is initially defined, and the regeneration system is active. Since the regeneration system is used in the same way under all conditions, the ATP-related trends observed in the early time interval are valid. However, we do not quantitatively interpret possible ATP depletion over a long timescale and do not base our results on this regime. The tug-of-war between contractility and crosslinking has been discussed and explored previously in great detail by Koenderink and coworkers suggesting a phase diagram in which fluid networks and those with local and global collapse exist (45). The emergence of maximal activity at comparably low ATP reveals a fundamental principle for active cytoskeletal networks: optimal non-equilibrium behavior requires both network compliance and motor force transmission. Networks that are too stiff resist motor-driven reorganization and motors that cycle too rapidly fail to coordinate their forces effectively. This finding has important implications for understanding cellular regulation of cytoskeletal dynamics. Cells operate in environments where ATP concentrations vary spatially and temporally depending on metabolic state, organelle proximity, and signaling activity (46–48). Our results suggest that local ATP levels could serve as a tunable parameter for modulating the balance between structural integrity and dynamic remodeling. The observed optimum at sub-millimolar ATP concentrations is particularly remarkable, as it falls within the range of physiologically relevant ATP fluctuations recently reported in cardiomyocytes (49), which contradicts the consensus of intracellular ATP levels being in the higher millimolar range (50). Moreover, our quantitative framework, which is linking ATP concentration to effective energy via MBR analysis, provides a novel approach for characterizing active matter systems that are not assessable via the ‘gold standard’ of using passive and active measurements at the same place with the same bead (33, 51).

**Figure 7.**
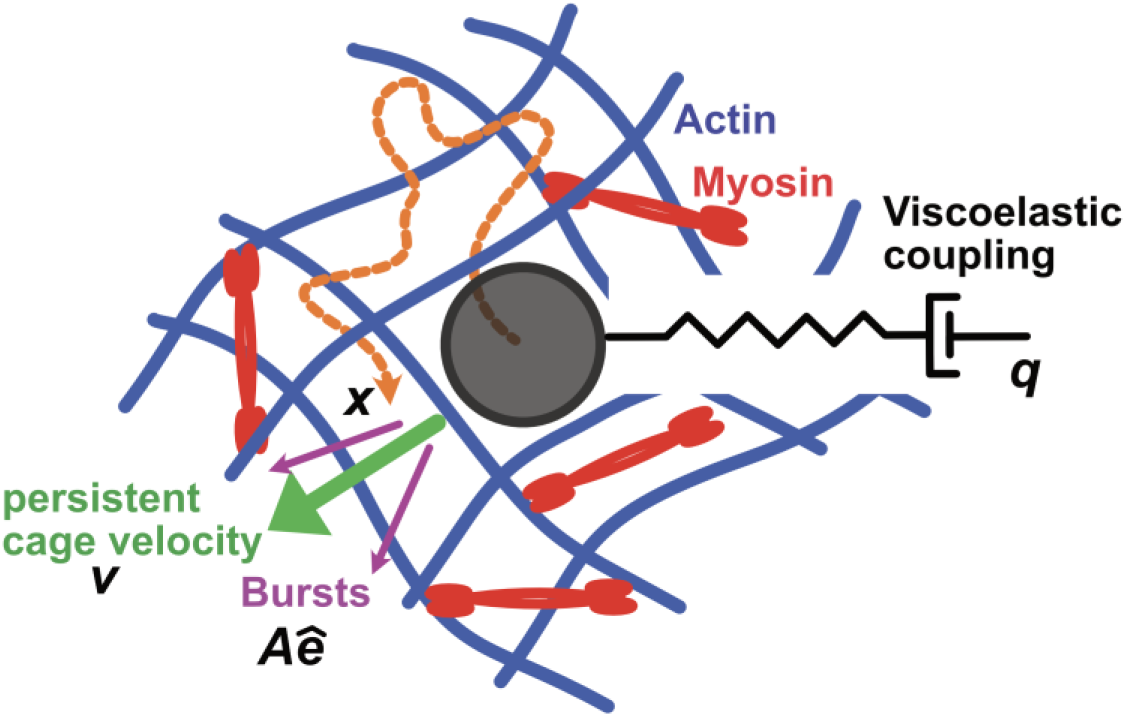
Schematic of the hidden-state active viscoelastic cage model. A tracer bead (black sphere) is embedded in an actomyosin network composed of actin filaments (blue) and myosin motor complexes (red). The bead trajectory (orange dashed) reflects a projection of a higher-dimensional hidden-state dynamics. The local actin cage exerts an elastic restoring force on the bead through a spring-dashpot element (right), representing the viscoelastic coupling between the observed bead position ***x*** and the hidden cage coordinate ***q***. Myosin-driven contractile events generate intermittent burst-like kicks (magenta arrows) to the hidden cage velocity, which in turn produce a persistent directed cage motion (green arrow, ***v***). This persistent velocity transports the local cage center ***q***, and the bead is dragged along via the elastic coupling *k*_*xq*_. The spring-dashpot element additionally captures delayed network relaxation through the hidden viscoelastic mode ***y***. In the passive thermal reference, myosin activity is absent (*r*_burst_ = 0) and the cage fluctuates only due to thermal noise. In the active condition, intermittent burst events representing coarse-grained myosin-driven rearrangements break fluctuation-dissipation balance and give rise to non-Gaussian bead displacements and altered back-relaxation statistics.

Future studies could extend this framework by investigating how other biochemical parameters such as myosin isoform, actin-binding protein composition, or filament length distribution modulate the ATP-dependent activity landscape. Additionally, computational models incorporating motor kinetics, network mechanics, and ATP-dependent feedback could provide mechanistic insights into the emergent scaling laws governing active cytoskeletal networks. In conclusion, our work demonstrates that cytoskeletal activity emerges from both motor kinetics and network mechanics. The identification of an optimal ATP concentration for maximizing non-equilibrium fluctuations reveals fundamental constraints on how cells might tune their internal dynamics through metabolic regulation, offering new perspectives on the biophysical principles underlying cellular self-organization.

## Materials and Methods

### Self-assembly of Actin Networks

Actin networks are formed by actin polymerization in the presence of myosin II and different ATP concentrations to study their effects on network properties. Lyophilized actin monomers (from rabbit skeletal muscle, Cytoskeleton, Denver, USA) and myosin (from rabbit skeletal muscle, Cytoskeleton, Denver, USA) are dissolved in deionized water. Actin monomers are diluted and stored in G-Actin Buffer (5 mM Tris HCl, 0.2 mM CaCl2, 0.2 mM ATP and 0.5 mM DTT; pH∼7.5) at 4 °C. For network formation, polymerization buffer (100 mM Tris HCl, 20 mM CaCl2, 500 mM KCl; pH∼7.5; 1:10 buffer:total sample volume), ATP (0 mM, 0.2 mM, 0.5 mM, 1 mM, 1.2 mM, 1.5 mM or 2 mM) and G-Actin buffer (for total sample volume of 10 μL) are mixed first. Next, the ATP regeneration system consisting of creatine phosphokinase (CPK, 75 units/mL, Sigma Aldrich, Taufkirchen, Germany) and phosphocreatine (CP, 1.25 mM, Sigma Aldrich, Taufkirchen, Germany) is added to keep the ATP levels constant over the measurement. Lastly, muscle actin is added to a concentration of 24 μM with 10 % being ATTO 488-labelled actin (from rabbit skeletal muscle, Hypermol, Bielefeld, Germany). Actin polymerization is induced by the high salt concentrations in the polymerization buffer. For actomyosin networks, 0.48 μM myosin II (from rabbit skeletal muscle, Cytoskeleton, Denver, USA) are added to the sample. For passive microrheology experiments, polystyrene beads are prepared as described in the following section.

### Passive Microrheology

Passive microrheology is used to quantify stiffness and activity of differently crosslinked actin networks (15, 18, 23, 51). Carboxylate-modified polystyrene beads with a diameter of 2 μm – large enough to avoid immediate incorporation into clusters yet small enough to sense motor-driven fluctuations (15) (Sigma-Aldrich, St. Louis, USA) - are passivated for 30 minutes in bovine serum albumin (BSA) to prevent binding to actin filaments or myosin motors and washed with G-Actin buffer to remove excess BSA. The particles are incorporated into the actin cytoskeletal networks, and their positions were monitored over a 180-second interval across samples of varying maturation times using a bright-field microscope equipped with a 60× objective (CFI Achromat FF, numerical aperture [NA] = 0.80, working distance = 0.3 mm; Nikon). Holographic illumination is achieved with a monochromatic light from a light-emitting diode (λ = 660 nm) and images are recorded at 138 fps from which beads were tracked using proprietary software (AFS, Lumicks BV, software: AFS-Tracking-G2-v1.1.5 (52)). The obtained single-particle tracking trajectories are detrended by a moving average filter. From the detrended trajectories, mean squared displacements (MSDs) (18), Van Hove correlations (18) and mean back relaxation (MBR) (20) are calculated using custom-written python codes. The van Hove distribution *P*(*Δx*; *τ*) was computed as the probability density of one-dimensional displacements *Δx*(*τ*) = *x*(*t* + *τ*) − *x*(*t*) at lag time *τ* for single exemplary trajectories.

The MSD was calculated as the time-averaged squared two-dimensional displacement ⟨Δ*r*(*τ*)^2^⟩ = ⟨Δ*x*(*τ*)^2^ + Δ*y*(*τ*)^2^⟩ over all lag times, pooled over all trials per condition. They were plotted up to *τ* = 10 s, where estimates remain statistically robust and ballistic flow doesn’t appear. The slope α between *τ* = 0.1 s and *τ* = 1 s was extracted from each individual MSD curve as a measure of bead diffusion.

From the averaged MSDs of each trial, *G*^′^(*ω*) and *G*^′′^(*ω*) were obtained using the Mason–Weitz method based on the generalized Stokes–Einstein relation (24, 25). The value of the storage modulus *G*^′^(*ω*) at approximately 0.1 Hz was taken as a measure of network stiffness. The effective energies *E*_0_/*k*_*B*_*T* were calculated from long-term MBR values according to Münker et al. (20).

### Confocal Laser Scanning Microscopy (CLSM Imaging)

Confocal laser scanning microscopy enables high-quality imaging of fluorescent samples in specific z-planes (53). A confocal laser scanning microscope (CLSM) composed of a confocal laser scanning unit (CLSM, FV4000, Olympus) on an inverted microscope (Olympus IX 83) equipped with a 60x water immersion objective (UPLSAPO60XW, NA = 1.2, Olympus) is employed to capture images for network structure comparison. Images are processed using FIJI software (54).

### Hidden-state active cage model

We simulated a two-dimensional overdamped tracer bead in a coarse-grained active viscoelastic network. For each Cartesian component, the hidden state was *Y*(*t*) = (*x*(*t*), *q*(*t*), *ν*(*t*), *y*(*t*))^⊤^, where *x* is the observed bead coordinate, *q* the local cage position, *ν* the persistent cage velocity, and *y* a delayed viscoelastic memory coordinate (44, 55). The bead was elastically coupled to both the moving cage and the memory coordinate, while the cage velocity generated persistent cage motion (55). Between active events, the dynamics were linear and Gaussian and were advanced exactly in discrete time. Writing d*Y* =*AY* d*t* + *K* d*W*, the update over one time step *Δt* was *Y*_*n*+1_ = *e*^*AΔt*^*Y*_*n*_ + *η*_*n*_, *η*_*n*_ ∼ 𝒩(0, *S*), with covariance *S* (see SI for details). Active actomyosin-like rearrangements were modeled as Poisson-distributed bursts acting on the hidden cage velocity *ν*. Each burst added an isotropic two-dimensional velocity kick to *ν*, causing the cage center to move persistently for a finite time. The bead was therefore not kicked directly but followed the actively moving cage through its elastic coupling. Passive reference simulations were obtained by setting the burst rate to zero while retaining thermal cage fluctuations. The simulation is similar to the one of Knotz et al. (43) and provides qualitatively the same results.

### Statistical Analyses

Statistical analyses were performed using two–sided Mann–Whitney U test (56). Significance levels were classified as follows: p<0.05 (*), p<0.01 (**), p<0.001 (***) and p<0.0001 (****). Data is presented as mean ± standard deviation or mean ± standard error as stated in the respective figure caption. The number of replicates and individual probes are listed in Table S1. P-values for statistical analyses are provided in Table S2-4.

## Acknowledgments

We gratefully acknowledge financial support from Deutsche Forschungsgemeinschaft (DFG, German Research Foundation: project-ID 449750155-RTG 2756, project A06).

