## Supplemental tables, figures and text. for "Myosin activity drives entangled actin networks out-of-equilibrium – a quantitative approach"

*Andreas Janshoff

### Supporting Information Text

#### Hidden-cage stochastic model

The following section describes the simulation procedure in greater detail. The hidden-state representation follows the standard Markovian embedding of generalized Langevin dynamics, in which exponentially decaying memory kernels are represented by auxiliary relaxation variables (2). These approaches replace non-Markovian memory kernels by auxiliary relaxation variables, yielding a Markovian dynamics in an enlarged hidden state space. Here, we use a distinct phenomenological construction: the bead itself is not self-propelled but is coupled to a hidden cage whose velocity receives intermittent active kicks. For each Cartesian coordinate, the system state is

$$Y(t) = \begin{pmatrix} x(t) \\ q(t) \\ v(t) \\ y(t) \end{pmatrix},$$

where $x(t)$ is the observed bead coordinate, $q(t)$ is the hidden cage center, $v(t)$ is the persistent cage velocity, and $y(t)$ is a hidden viscoelastic memory coordinate. The two Cartesian components are simulated independently with identical scalar dynamics.

The bead is assumed to be overdamped and coupled elastically to both the cage center and the viscoelastic memory coordinate:

$$\mathrm{d}x = \frac{-k_{xq}(x-q) - k_{xy}(x-y)}{\gamma_x}\,\mathrm{d}t + \sqrt{\frac{2k_\mathrm{B}T_x}{\gamma_x}}\,\mathrm{d}W_x.$$

The first elastic term couples the bead to the active cage center, while the second term couples it to the hidden viscoelastic memory mode.

The hidden cage center evolves according to

$$\mathrm{d}q = \left(v - \lambda_q q\right)\mathrm{d}t + \sqrt{2D_q}\,\mathrm{d}W_q,$$

Activity is introduced by Poisson-distributed burst events acting on the hidden cage velocity. During a time step $\Delta t$, the number of bursts is sampled from

$$N_\text{burst} \sim \mathrm{Poisson}(\lambda_\text{burst}\,\Delta t),$$

where $\lambda_\text{burst}$ is the burst rate. Each burst adds an isotropically oriented kick to the cage velocity,

$$v \longrightarrow v + \kappa.$$

In two dimensions, the kick direction is sampled uniformly on the circle, and the kick magnitude is sampled from the prescribed amplitude distribution.

Importantly, active bursts are not applied directly to the observed bead coordinate. Instead, they perturb the hidden cage velocity. The cage center then moves persistently, and the bead follows through the elastic coupling $k_{xq}$. This implements active rearrangements of the local environment rather than direct active forcing of the bead. Between active burst events, the hidden-state dynamics are linear and Gaussian (this is also visible in the corresponding van Hove plots, see main text). For one Cartesian component, e.g. $x(t)$, we obtain

$$\mathrm{d}Y(t) = AY(t)\mathrm{d}t + K\,\mathrm{d}W(t),$$

The drift matrix *A* is

$$A = \begin{pmatrix} -\dfrac{k_{xq}+k_{xy}}{\gamma_x} & \dfrac{k_{xq}}{\gamma_x} & 0 & \dfrac{k_{xy}}{\gamma_x} \\ 0 & -\lambda_q & 1 & 0 \\ 0 & 0 & -\dfrac{1}{\tau_v} & 0 \\ \dfrac{k_{yx}}{\gamma_y} & 0 & 0 & -\dfrac{k_{yx}}{\gamma_y} \end{pmatrix},$$

and the noise matrix *K* is

$$K = \begin{pmatrix} \sqrt{\dfrac{2k_{\mathrm{B}}T_x}{\gamma_x}} & 0 & 0 & 0 \\ 0 & \sqrt{2D_q} & 0 & 0 \\ 0 & 0 & \sqrt{\dfrac{2D_v}{\tau_v}} & 0 \\ 0 & 0 & 0 & \sqrt{\dfrac{2k_{\mathrm{B}}T_y}{\gamma_y}} \end{pmatrix}.$$

The linear Gaussian part is advanced exactly over each time step. The exact discrete-time transition is

$$Y_{n+1} = BY_n + \eta_n, B = e^{A\Delta t}$$

where $\eta_n$ is a zero-mean Gaussian random vector with covariance

$$S = \int_0^{\Delta t} e^{Au} K K^{\top} e^{A^{\top}u}\,\mathrm{d}u$$

The covariance matrix $S$ was computed using the Van Loan block-matrix method. Gaussian increments with covariance $S$ were generated by a Cholesky or eigenvalue-based square-root factorization. This exact update avoids time-discretization errors associated with an Euler-Maruyama approximation of the linear part. Active burst jumps are applied before the exact linear Gaussian propagation over each time step.

All simulations were performed in dimensionless units. The model is intended as a minimal coarse-grained active hidden-cage model rather than as a directly calibrated representation of a specific experimental system. We therefore use simulation units in which the bead friction coefficient and the bead thermal energy scale are set to unity,

$$\gamma_x = 1, k_{\mathrm{B}}T_x = 1.$$

With this convention, the free thermal diffusion coefficient of the observed bead is

$$D_x = \frac{k_{\mathrm{B}}T_x}{\gamma_x} = 1$$

Thus, times, lengths, stiffnesses, and diffusion coefficients are reported in the corresponding nondimensional simulation units. More generally, dimensional variables could be recovered by introducing a reference length scale $L_0$, a reference time scale $T_0$, and a reference energy scale $E_0$. Dimensionless variables are then defined by

$$\tilde{x} = \frac{x}{L_0}, \tilde{q} = \frac{q}{L_0}, \tilde{y} = \frac{y}{L_0}, \tilde{t} = \frac{t}{T_0}, \tilde{v} = \frac{T_0}{L_0} v$$

The corresponding dimensionless stiffnesses, frictions, and diffusion coefficients are

$$\tilde{k} = \frac{kL_0^2}{E_0}, \tilde{\gamma} = \frac{\gamma L_0^2}{E_0 T_0}, \tilde{D} = \frac{DT_0}{L_0^2}$$

In the simulations below, all tildes are dropped for notational simplicity. The quantity denoted by $k_\mathrm{B}T_x$ should therefore be understood as the thermal energy scale of the bead bath in simulation units. Similarly, $k_\mathrm{B}T_y$ sets the noise scale of the hidden viscoelastic mode. In contrast, the active burst process is not thermal: burst events inject nonequilibrium kicks into the hidden cage velocity and are not constrained by a fluctuation-dissipation relation. Consequently, no single equilibrium temperature can be assigned to the full active system.

**Dependency of the MBR on the environment**

The mean back relaxation (MBR) is sensitive to whether a selected large displacement is followed by a restoring motion or by continued motion in the same direction. For a selected displacement

$$d = x(s) - x(s - \tau)$$

the MBR is defined as (see main text)

$$\mathrm{MBR}(\tau, t, l) = \left\langle -\frac{x(s + t) - x(s)}{x(s) - x(s - \tau)} \middle| |x(s) - x(s - \tau)| > l \right\rangle.$$

Thus, positive MBR values indicate back-relaxation, i.e., the bead tends to move opposite to the selected displacement (hence, the term back relaxation). Negative values indicate persistence, which means that the bead continues to move in the same direction as the selected displacement. In the active cage model, two competing mechanisms control the MBR. First, active bursts drive the hidden cage velocity $v$ and therefore generate persistent cage motion. Second, the viscoelastic hidden coordinate $y$ stores elastic memory and tends to pull the bead back. The observed MBR is therefore determined by the balance between active persistence and viscoelastic restoring forces.

The parameter $k_{xy}$ controls the strength of the coupling between the observed bead coordinate $x$ and the hidden viscoelastic coordinate $y$. Increasing $k_{xy}$ strengthens the restoring force:

$$F_{xy} = -k_{xy}(x - y).$$

Therefore, when the bead is displaced by active cage motion, a larger $k_{xy}$ stores more elastic deformation in the hidden viscoelastic mode and pulls the bead back more strongly. This suppresses persistent forward motion and increases the MBR value at a given $t = \tau$. To illustrate this, we have chosen a reasonable lag time and plotted the MBR value as a function of $k_{xy}$ (Figure S5). This plot shows that the stronger the coupling to the medium is (the stiffer the network), a less pronounced persistence of bead movement can be found (lower MBR). This is intuitively clear as in a very stiff network any movement is stalled. It stresses the importance of comparing equally stiff networks when it comes to back relaxation.

The parameter $k_{xq}$ controls the strength of the coupling between the bead and the active cage center $q$. The corresponding force is

$$F_{xq} = -k_{xq}(x - q).$$

Although this term has the form of an elastic coupling, in the active model $q$ is not a fixed passive trap. Instead, $q$ is driven by the hidden persistent cage velocity $v$, which receives active burst kicks. Therefore, increasing $k_{xq}$ makes the bead follow the active cage more strongly (Figure S6). Therefore, the MBR drops and eventually becomes negative (Figure S6).

The burst-rate parameter $\lambda$ controls how often active kicks are applied to the hidden cage velocity $v$. Increasing $\lambda$ increases the frequency of active rearrangement events. At fixed burst amplitude, this increases the probability that a selected large displacement is associated with recent active cage motion. The bead then continues moving in the same direction after the selected displacement, causing the MBR to decrease (Figure S7). The viscoelastic damping parameter $\gamma_y$ has a different effect. The hidden viscoelastic coordinate obeys

$$\mathrm{d}y = -\frac{k_{yx}}{\gamma_y}(y - x)\mathrm{d}t + \sqrt{\frac{2k_\mathrm{B}T_y}{\gamma_y}}\ \mathrm{d}W_y.$$

The corresponding relaxation time is approximately:

$$\tau_y = \frac{\gamma_y}{k_{yx}}.$$

Therefore, increasing $\gamma_y$ makes the viscoelastic coordinate slower (Figure S7). On the observation timescale $t \sim \tau = 1$, a larger $\gamma_y$ means that $y$ responds more slowly to bead motion and acts more like an anchor. This increases stored elastic memory and promotes back-relaxation (Figure S7).

### Figures

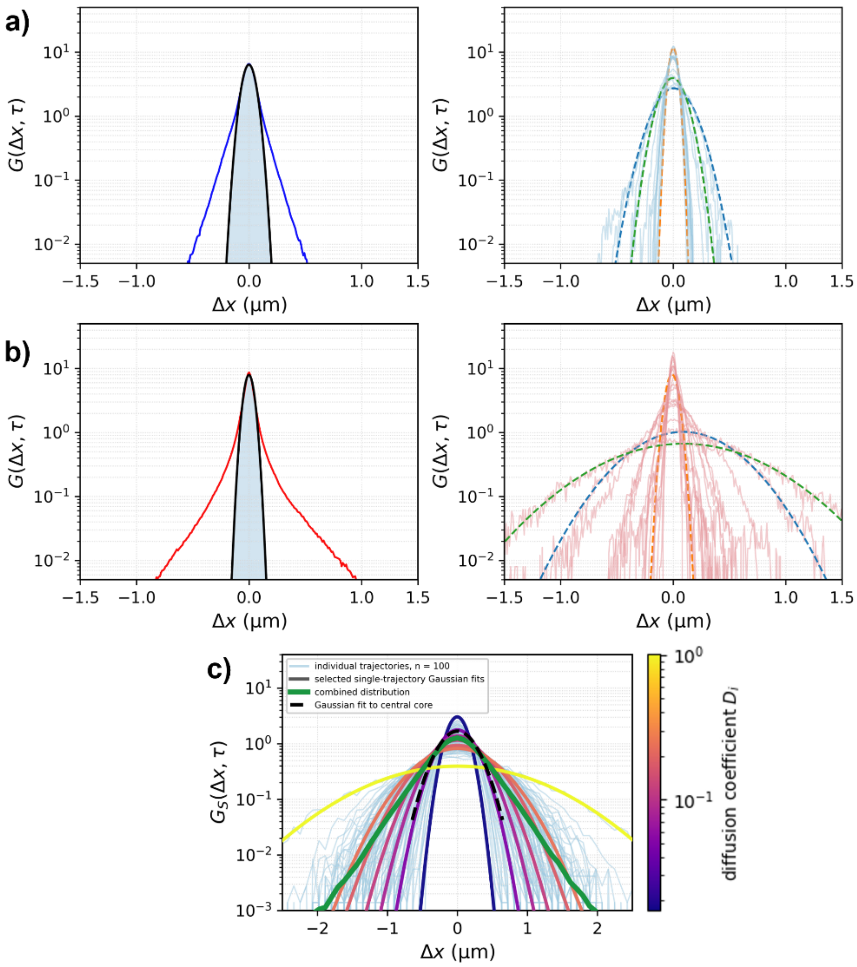


**Fig. S1.** Van Hove correlation functions computed from the x-dimension of two-dimensional bead trajectories of probe particles. a) On the left, combined Van Hove distribution calculated by pooling all displacements from all trajectories of beads embedded in actin networks with 0.5 mM ATP at a sample age of 20 minutes are shown. The blue line represents the experimental data, and the corresponding fit is shown in black. On the right, van Hove distributions a set of randomly chosen single trajectories of that condition with three exemplary Gaussian fits illustrate how the ensemble curve is composed of gaussians with different widths. b) Similar to a, but for actomyosin networks with 0.5 mM ATP. Here, the van Hove distributions for individual trajectories as well as the pooled data are depicted in red. Actin concentration: 24 µM, myosin concentration: 0.48 µM, $\tau$: 1s. c) Illustration of heterogeneity-induced non-Gaussian tails in the self-part of the van Hove distribution derived from simulated data. Each trajectory was generated with a fixed trajectory-specific diffusion coefficient $D_i$, drawn from a log-normal distribution. For a given trajectory $i$, the displacements are therefore purely Gaussian. The light-blue curves show the empirical displacement distributions of the individual trajectories. The colored solid curves show selected single-trajectory Gaussian fits; their color encodes the corresponding diffusion coefficient $D_i$ on a logarithmic scale, as indicated by the color bar on the right side. Narrow curves correspond to low-mobility trajectories with small $D_i$, whereas broader curves correspond to high-mobility trajectories with large $D_i$. Although each trajectory is Gaussian, the combined distribution (solid green) shows clear heavy tails on the semi-log scale. This arises because the overall distribution is a superposition of Gaussians with different variances. The black dashed line is a Gaussian fit to the central part only: it captures the narrow peak around Δx = 0 but severely underestimates large displacements. The deviation of the green curve from this central fit shows that the observed tails do not require non-Gaussian single-trajectory dynamics—they can result solely from heterogeneity in diffusion coefficients.

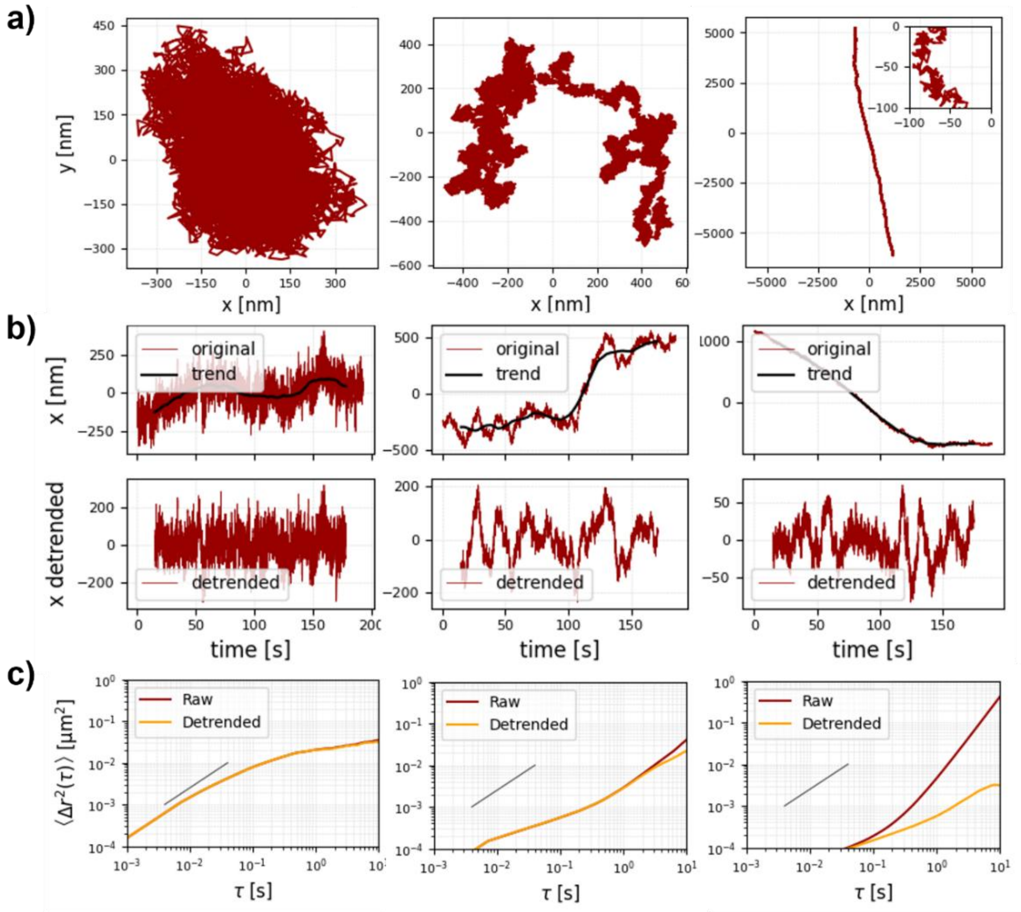


**Fig. S2.** Detrending of exemplary trajectories of beads embedded in actomyosin networks. a) Different types of 2D trajectories of beads embedded in actomyosin networks. b) Illustration of the detrending process by subtraction of the general trend depicted in black from the one-dimensional raw trajectory (upper red curve) leading to the detrended trajectory shown below. The trend was defined by applying a moving average filter with a kernel size of 4001 datapoints. c) Mean squared displacements calculated from the respective raw (red) and detrended trajectories (orange).

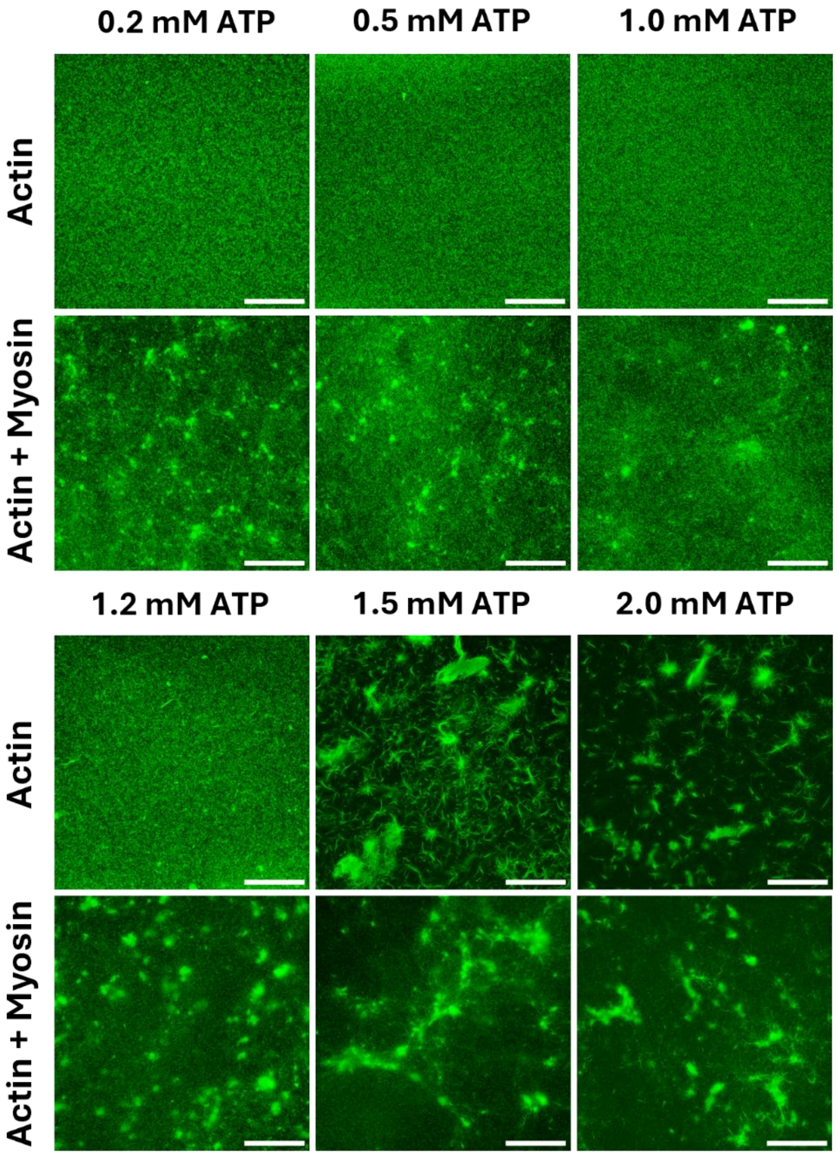


**Fig. S3.** Confocal fluorescence images of actin and actomyosin networks polymerized at all tested ATP concentrations. The images were taken 20 minutes after sample preparation. Histogram equalization was applied to enhance contrast in Fiji (1). Actin concentration: 24 μM with 10% being ATTO-488 labelled actin; myosin concentration: 0.48 μM. The scale bars represent 50 μm.

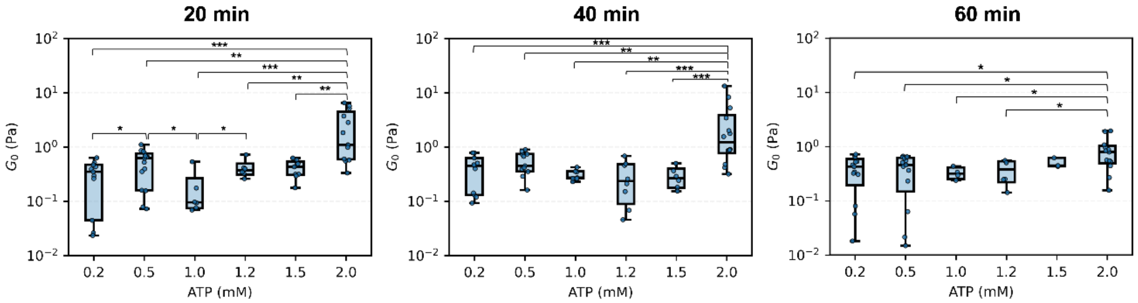


**Fig. S4.** Plateau moduli $\boldsymbol{G_0}$ (approx. obtained at 0.1 Hz) calculated from MSDs of actin and actomyosin networks polymerized with different concentrations of ATP. The plots represent results from measurement at 20, 40 and 60 minutes of sample age. Actin concentration: 24 μM. All pairwise comparisons between conditions were performed using two-sided Mann–Whitney U test. Significance levels were indicated as $p<0.05$ (*), $p<0.01$ (**), $p<0.001$ (***). All p-values and sample statistics can be found in Table S1 and Table S4.

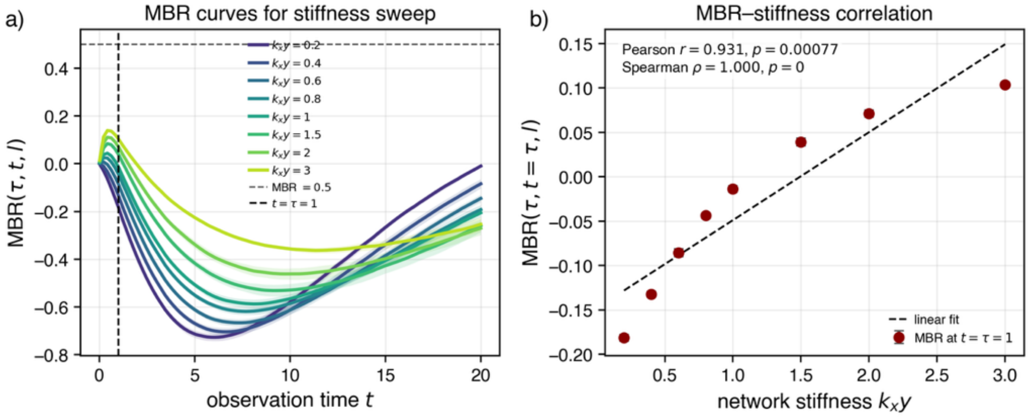


**Fig. S5.** Dependence of the mean back relaxation on the viscoelastic coupling stiffness $k_{xy}$. (a) MBR curves for different values of $k_{xy}$. The vertical dashed line marks the evaluation time $t = \tau = 1$, and the horizontal dashed line indicates the passive long-time reference value MBR $= 0.5$. Increasing $k_{xy}$ shifts the MBR upward because the bead is more strongly coupled to the viscoelastic memory coordinate $y$. (b) Correlation between $k_{xy}$ and $\mathrm{MBR}(\tau, t = \tau, l)$. The correlation shows that stronger viscoelastic coupling produces stronger back-relaxation and reduces active forward persistence.

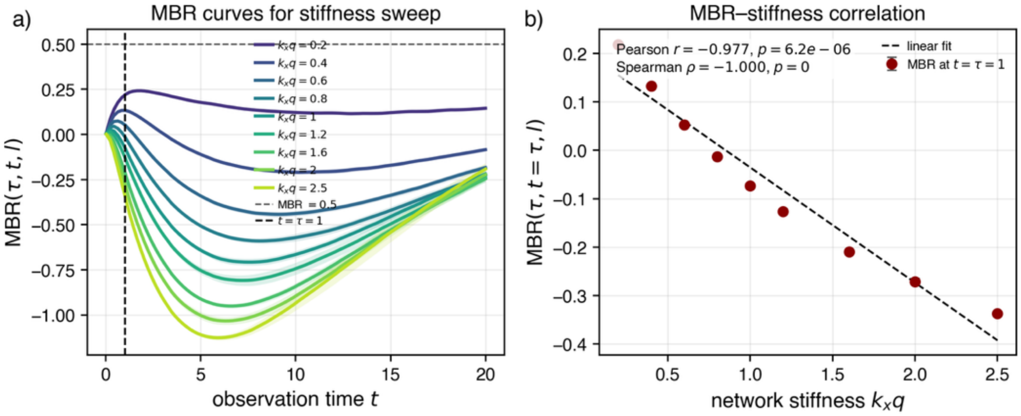


**Fig. S6.** Dependence of the mean back relaxation on the bead-cage coupling stiffness $k_{xq}$. (a) MBR curves for different values of $k_{xq}$. The vertical dashed line marks $t = \tau = 1$. For small $k_{xq}$, the bead is only weakly coupled to the active cage and the MBR can remain positive. For larger $k_{xq}$, the bead follows the active cage more strongly and the intermediate-time MBR becomes negative. (b) Correlation between $k_{xq}$ and $\mathrm{MBR}(\tau, t = \tau, l)$. The strong negative correlation shows that stronger coupling to the active cage enhances persistent motion and therefore lowers the MBR.

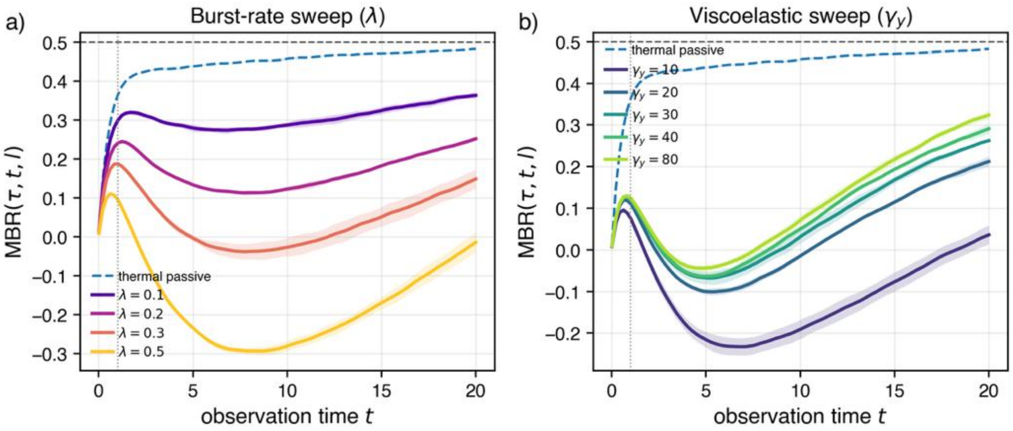


**Fig. S7.** Effect of active burst rate and viscoelastic damping on the MBR. (a) Burst-rate sweep. Increasing the burst rate $\lambda$ increases the frequency of active kicks applied to the hidden cage velocity $v$. This enhances persistent active motion and lowers the MBR at intermediate times. (b) Viscoelastic damping sweep. Increasing $\gamma_y$ increases the relaxation time $\tau_y = \gamma_y / k_{yx}$ of the hidden viscoelastic coordinate $y$. The hidden network coordinate then responds more slowly and behaves more like an elastic anchor on the observation timescale. This enhances back-relaxation and shifts the MBR upward. The dashed blue curve shows the thermal passive reference, the horizontal dashed line marks MBR $= 0.5$, and the vertical dotted line marks $\tau = 1$.

**Tables**

**Table S1.** Statistical information on passive microrheology measurements.

| ATP concentration | + Myosin | Measurements | Probes |
|---|---|---|---|
| 0.2 mM | No | 15 | 222 |
| | Yes | 16 | 165 |
| 0.5 mM | No | 17 | 230 |
| | Yes | 25 | 272 |
| 1 mM | No | 9 | 123 |
| | Yes | 19 | 225 |
| 1.2 mM | No | 8 | 77 |
| | Yes | 19 | 210 |
| 1.5 mM | No | 11 | 111 |
| | Yes | 18 | 275 |
| 2 mM | No | 17 | 169 |
| | Yes | 16 | 181 |

**Table S2.** Measurement statistics of Figure 4 and Figure S3. Ratio of the slopes and plateau heights of the ensemble-averaged MSDs for actomyosin networks and the corresponding networks without myosin, the respective propagated errors and p-values. The uncertainty of the ratio was calculated using standard Gaussian error propagation, combining the relative standard errors of both measurements in quadrature. Significance was assessed using a two-sided Mann–Whitney U test comparing MSD plateau heights for actomyosin networks and the respective networks without myosin.

| | **Scaling exponent α** | | | | |
|---|---|---|---|---|---|
| **ATP** | **Mean_Actin** | **Mean_Myo** | **Ratio** | **Error** | **p-value** |
| **0.2 mM** | 0.349 | 0.638 | 1.827 | 0.072 | 6.142e-28 |
| **0.5 mM** | 0.381 | 0.641 | 1.683 | 0.054 | 3.696e-26 |
| **1 mM** | 0.405 | 0.393 | 0.971 | 0.035 | 0.0007999 |
| **1.2 mM** | 0.345 | 0.546 | 1.584 | 0.059 | 7.27e-10 |
| **1.5 mM** | 0.318 | 0.375 | 1.181 | 0.063 | 0.04896 |
| **2 mM** | 0.217 | 0.347 | 1.596 | 0.130 | 1.496e-05 |

**Table S3**. p-values of Figure 5 for pairwise comparisons between all conditions for $E_0/k_BT$ r values measured at 20 minutes of sample age. Statistical analyses were performed using the two-sided Mann–Whitney U test.

| | Effective energies $E_0/k_BT$ | | | | |
|---|---|---|---|---|---|
| **ATP** | **Mean_Actin** | **Mean_Myo** | **SEM_Actin** | **SEM_Myo** | **p-value** |
| **0.2 mM** | 0.054 | 0.438 | 0.003 | 0.031 | 1.212e-49 |
| **0.5 mM** | 0.065 | 0.424 | 0.003 | 0.031 | 3.356e-35 |
| **1 mM** | 0.028 | 0.116 | 0.002 | 0.010 | 3.198e-17 |
| **1.2 mM** | 0.061 | 0.228 | 0.003 | 0.019 | 2.533e-08 |
| **1.5 mM** | 0.051 | 0.097 | 0.005 | 0.008 | 9.531e-02 |
| **2 mM** | 0.043 | 0.118 | 0.006 | 0.013 | 5.308e-05 |

**Table S4.** p-values of Figure S4 for pairwise comparisons between all conditions measured at 20, 40, and 60 minutes after sample preparation. Statistical analyses were performed using the two-sided Mann–Whitney U test.

| **20 minutes** | | | | | |
|---|---|---|---|---|---|
| **Condition** | **0.5 mM** | **1 mM** | **1.2 mM** | **1.5 mM** | **2 mM** |
| **0.2 mM** | 4.455e-02 | 8.044e-01 | 5.002e-01 | 3.247e-01 | 4.636e-04 |
| **0.5 mM** | | 4.947e-02 | 7.108e-01 | 3.468e-01 | 8.208e-03 |
| **1 mM** | | | 4.988e-02 | 5.062e-02 | 1.836e-04 |
| **1.2 mM** | | | | 9.678e-01 | 4.169e-03 |
| **1.5 mM** | | | | | 2.198e-03 |
| **40 minutes** | | | | | |
| **Condition** | **0.5 mM** | **1 mM** | **1.2 mM** | **1.5 mM** | **2 mM** |
| **0.2 mM** | 3.560e-01 | 4.626e-01 | 4.757e-01 | 6.452e-01 | 9.300e-04 |
| **0.5 mM** | | 5.280e-02 | 8.811e-02 | 7.560e-02 | 7.976e-03 |
| **1 mM** | | | 5.956e-01 | 8.884e-01 | 1.381e-03 |
| **1.2 mM** | | | | 6.965e-01 | 9.876e-04 |
| **1.5 mM** | | | | | 7.016e-04 |
| **60 minutes** | | | | | |
| **Condition** | **0.5 mM** | **1 mM** | **1.2 mM** | **1.5 mM** | **2 mM** |
| **0.2 mM** | 9.669e-01 | 4.702e-01 | 8.746e-01 | 4.973e-01 | 1.440e-02 |
| **0.5 mM** | | 5.187e-01 | 8.464e-01 | 5.531e-01 | 2.253e-02 |
| **1 mM** | | | 7.546e-01 | 8.225e-02 | 2.926e-02 |
| **1.2 mM** | | | | 3.543e-01 | 3.372e-02 |
| **1.5 mM** | | | | | 2.661e-01 |
